# Interaction between β-lactoglobulin and EGCG under high-pressure by molecular dynamics simulation

**DOI:** 10.1101/2021.08.09.455733

**Authors:** Yechuan Huang, Xicai Zhang, Huayi Suo

## Abstract

The binding between β-lactoglobulin and epigallocatechin gallate (EGCG) under the pressure of 600 MPa was explored using molecular docking and molecular dynamics (MD) simulation. EGCG bound mainly in two regions with site 1 in internal cavity of the β-barrel and site 2 on the surface of protein. 150 ns MD was performed starting from the structure with the lowest binding energy at the two sites in molecular docking, respectively, it was found that the system fluctuated greatly when small molecule bound to site 2 at 0.1 MPa. The protein fluctuation and solvent accessible surface area became smaller under high-pressure. The binding of small molecules made the protein structure more stable with increasing of α-helix and β-sheet, while high-pressure destroyed α-helix of protein. The binding free energy of small molecules at site 1was higher than that at site 2 under 0.1 MPa, with higher van der Waals and hydrophobic interaction at site 1 while more hydrogen bonds at site 2. The binding free energy of both sites decreased under high-pressure, especially at site 1, causing binding free energy at site 1 was lower than that at site 2 under high-pressure.

## 1. Introduction

Milk is an essential food for modern people. In addition to direct drinking, there are also many milk drinks, for example, tea was added into milk to make the product more diverse in form and taste and more comprehensive in nutrition (Stanner, 2007). Tea contains a lot of polyphenols, which have many physiological functions, such as antioxidation, reducing the incidence of cancer and cardiovascular (Carocho & Ferreira, 2013). Polyphenols are easy to form complex with milk protein, such as α-and β-caseins (Hasni et al., 2011), β-lactoglobulin (Kanakis et al., 2011), thus affecting the quality of products and the biological activity of polyphenols. Many scholars have studied the binding of protein with polyphenols, such as the binding type, binding site and binding mechanism (Li et al, 2018; Zhang, Wang, & Pan, 2012; Gholami & Bordbar, 2014). The binding between them is mainly non covalent, such as van der Waals force, hydrophobic interaction, electrostatic interaction and hydrogen bond, and covalent cross-linking also occurs under certain conditions (Jakobek, 2015; Sui et al., 2018).

At present, dairy products are still mainly sterilized by heat, thus having some negative effect on product quality. Ultra-high pressure is a kind of cold sterilization technology, which has good sterilization effect without destroying the flavor and nutrient of small molecules in food. Therefore, it is widely used in food, including dairy products (Stratakos et al., 2019; Malinowska-Pańczyk, 2020). The application of ultra-high pressure in milk will change the structure of macromolecular milk protein, and then affect the quality of dairy products. Some researchers reported the effect of high-pressure treatment on the structure of milk protein (Russo et al., 2013; Considine, Patel, Singh, & Creame, 2007). In terms of the effect of ultra-high pressure on the binding of polyphenols and proteins, Chen, Wang, Feng, Jiang, and Miao (2019) reported that soybean protein and tea polyphenols were mainly crosslinked by hydrophobic interaction and hydrogen bond under 400 MPa, protein had protective effect on the biological activity of tea polyphenols, and the addition of tea polyphenols could also reduce the change of protein structure under high pressure. To our best understanding, there was no report on how high pressure affected the binding between polyphenols and proteins in milk drinks.

Spectroscopy, calorimetry and atomic force microscopy are widely used to study protein structure, but only static structure is measured. Protein in organism is always in dynamic change. Now molecular dynamics simulation has become another means with enough small time and space scale to understand protein structure from molecular level after experimental and theoretical means (Chen, 2017). The binding process and mechanism of milk protein with many small molecules were studied by molecular modelling, such as citrus flavonoids (Sahihi & Ghayeb, 2014), curcumin (Mohammadi, Sahihi, & Khalegh Bordbar, 2015), apigenin (Zhu, Li, Wu, Li, & Sun, 2020), capsaicin (Zhan et al., 2020), and tea polyphenols (Kanakis et al, 2011). Furthermore, a few scholars have simulated the structural changes of some proteins (Kurpiewska, Miłaczewska, & Lewiński, 2019; Sarupria, Ghosh, García, & Garde, 2010; Kitchen, Reed, & Levy, 1992) or enzymes (Chen, 2017; Smolin & Winter, 2006) under high pressure. However, the simulation of the interaction between small molecules and proteins under high pressure has not been reported so far.

β-lactoglobulin is rich in milk with a lot of genetic variants particularly A and B phenotypes (Sawyer & Kontopidis, 2000), because of good function and nutritional characteristics, and good affinity for both hydrophobic and amphoteric ligands, it is widely used in food industry. Each monomer of β-lactoglobulin contains 162 amino acids with a molecular weight of 18.4 kDa (Brownlow et al, 1997).

As shown in Fig. 1, it has a calyx composed of 8-standard antiparallel β-sheets and an α-helix structure on the outside. EGCG, namely epigallocatechin gallate, is one of the most important components in tea polyphenols, which has very important health care function for human body (Yan, Zhong, Duan, Chen, & Li, 2020; Lecumberri, Dupertuis, Miralbell, & Pichard, 2013). Therefore, molecular dynamics simulation was used to study the changes of binding energy and binding mechanism between β-lactoglobulin and EGCG at 600 MPa in this study, so as to lay a theoretical foundation for further improving the quality of tea milk beverage and the application of high-pressure technology in milk beverage.

**Fig.1.**
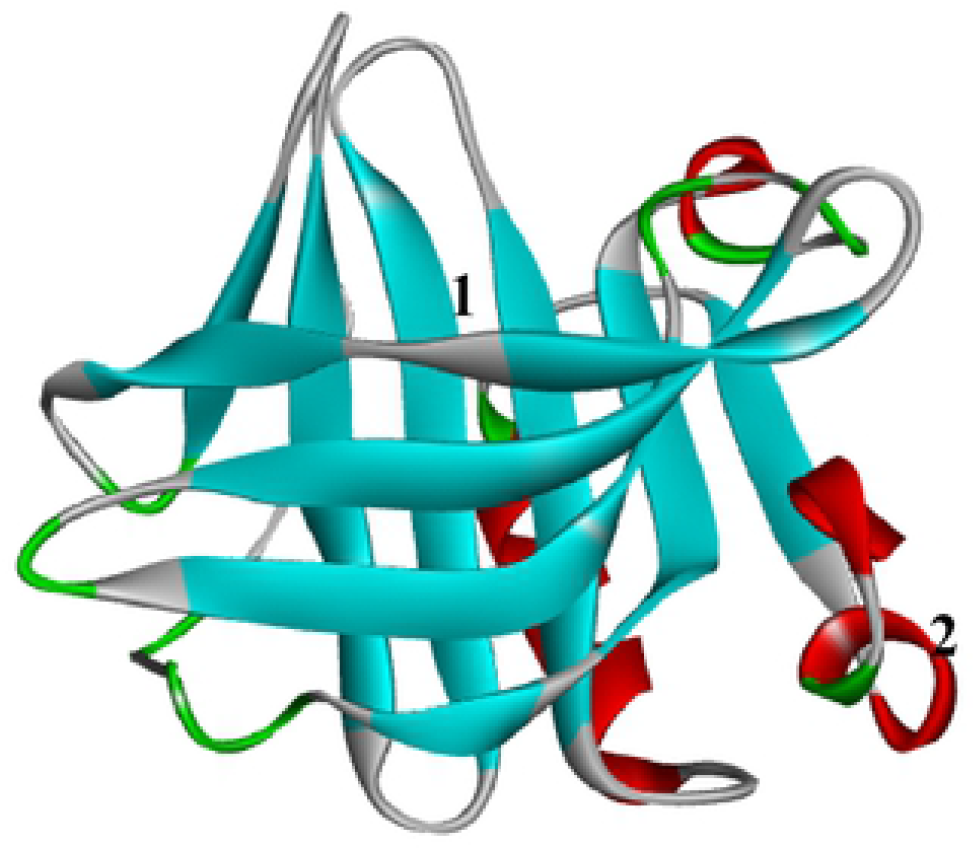
The main binding sites of EGCG β-laetoglobulin.

## 2. Materials and methods

### 2.1 Preparation of ligands and protein

β-lactoglobulin was downloaded from RCSB website (Rose et al., 2011) with an ID of 3npo, which had no missing atom, no crystallized ligand and had a reasonable resolution. Then the crystal water of protein was removed with PyMOL software, and the Gastieger charges and missing hydrogen atoms were added to protein by AutoDock Tools (version 1.5.6). EGCG was prepared by ChemDraw Professional 17.1 and Chem3D 17.1, and the structure was optimized by the self-contained MM2 force field. The coordinate files (PDBQT) of EGCG and β-lactoglobulin were prepared in AutoDock Tools environment respectively.

### 2.2 Molecular docking of β-lactoglobulin with EGCG

To investigate the binding sites of EGCG to β-lactoglobulin, a blind docking was performed using AutoDock Vina program (Trott & Olson, 2010) to search the top 20 optimal conformations. In the process of docking, the small molecule was set to be fully flexible and the center coordinate was the geometric center of the receptor, a box of 40 * 40 * 40 Å^3^ was set with a grid spacing of 1 Å, and the docking parameters of energy_ range and exhaustiveness were set to 5 and 100, respectively.

### 2.3 Molecular dynamics simulation of EGCG binding to β-lactoglobulin under high pressure

Although molecular docking can explore the binding site and binding energy between small molecule and protein, they could not fully understand the dynamic changes in the binding process. Therefore, 150 ns of molecular dynamics simulation was conducted. The optimal conformations of site 1 and site 2 derived from docking simulations were selected as the initial structure to perform molecular dynamics simulation with GROMACS (2019.6) software package (Abraham et al., 2015; Van Der Spoel et al., 2005). The topology parameters of protein and small molecule were generated by Gromacs program and Automated Topology Builder (ATB version 3.0) server (Koziara, Stroet, Malde, & Mark, 2014), respectively. The GROMOS54a7 force field was applied to describe the interaction (Schmid et al, 2011), and the complex was immersed into a periodic water box of cube shape using a single point charge (SPC) water model. The shortest distance between the protein and the edge of the box was 1 nm, and Na^+^ was added to make the system electrically neutral. Energy minimization of the system was performed to relieve unfavorable interactions using the steepest descent method (Hess, Kutzner, van der Spoel, & Lindahl, 2008). Simulation comprised of equilibration and production phases, firstly a 400 ps simulation in NVT ensemble was carried out to equilibrate the system, and temperature was maintained at 300 K using velocity-rescale method. Then another 400 ps equilibrium in NPT ensemble was followed using the Parrinello–Rahman barostat to maintain the pressure at 0.1 or 600 MPa. Finally, a 150 ns MD was performed with a time step of 2 fs. PME (Particle Mesh Ewald) was used to calculate long-range electrostatics, and the cutoff of van der Waals and Coulomb interaction was set to 1.0 nm (Chen, 2017; Zhan et al, 2020; Sahihi & Ghayeb, 2014; Gholami & Bordbar, 2014). The atomic coordinates were recorded to the trajectory file every 5 ps.

At the end of the simulation, the periodic boundary conditions were removed, and then the corresponding commands of GROMACS analysis tools was used to evaluate the root mean square deviation (RMSD), root mean square fluctuation (RMSF), gyration radius (Rg), hydrogen bond between proteins, protein secondary structure and solvent accessible surface area. Origin 8.0 software was used for plotting.

### 2.3 Binding free energy between small molecule and protein using MMPBSA

The 500 snapshot structures of each complex were extracted from the trajectory file of last 50 ns at an interval of 100 ps, and then MM-PBSA (molecular mechanics Poisson – Boltzmann surface area) method, which was implemented in g_mmpbsa tool, was used to calculate the binding free energy between small molecule and protein (Kumari, Kumar, & Lynn, 2014). Solvent accessible surface area (SASA) was used as a non-polar solvent model.

### 2.4 The 2-D plot and surface structural for interaction between EGCG and β-lactoglobulin

The average PDB structure of the complex from 100ns to 150ns under different simulation conditions was extracted, and the hydrophobic and hydrogen bonding interactions between small molecule and protein at the binding sites were analyzed using LigPlot+ (V 2.2) software (Laskowski & Swindells, 2011). The surface structure of the protein and its binding to small molecule was drawn by PyMOL.

## 3. Results and discussion

### 3.1 The binding sites of EGCG to β-lactoglobulin

Through molecular docking, it was found that the top 20 high energy binding sites of EGCG in β-lactoglobulin were mainly distributed in two regions, as shown in Fig. 1, 14 of which were at the top of the hydrophobic cavity (site 1), and another 6 were located in the protein surface (site 2). The average docking energy in site 1 was greater than that in site 2. There were many reports on the binding sites of polyphenols to β-lactoglobulin, most of which believed that small molecules can bound to the hydrophobic cavity and protein surface simultaneously, and the binding energy in hydrophobic cavity was greater (Al-Shabib et al, 2018; Dan et al., 2019). However, Gholami and Bordbar (2014), and Li et al (2018) reported that the small molecules were adsorbed only on the surface of β-lactoglobulin. These results may be related to the differences in the structure of small molecules, the initial structure of docking protein or the simulation conditions.

### 3.2 Molecular dynamics simulation of EGCG binding to β-lactoglobulin

#### 3.2.1 Changes of RMSD in molecular dynamics simulation

RMSD (root mean square deviation) is a measure of the average deviation of protein structure from the original conformation at a given time and is an important indicator to evaluate the stability of the research system. Figure 2 showed the RMSD comparison of protein (a) and complex (b) under different treatment conditions. It could be seen from Fig.2 (a) that the protein fluctuated greatly when EGCG bound to site 2 at 0.1 MPa, and reached equilibrium after 80 ns, while the other three systems achieved equilibrium at about 50 ns. The protein average RMSD after 100ns was 0.2898nm when EGCG bound to site 2 at 0.1MPa, which was significantly higher than that for site 1 at 0.1MPa (0.2459nm), site 1 at 600 MPa (0.2464nm) and site 2 at 600 MPa (0.2513nm), indicating that the protein structure would fluctuate more when the small molecule was bound to site 2 under normal pressure, but the fluctuation would decrease after high pressure treatment. It could be seen from Fig.2 (b) that the RMSD fluctuation of the complex was consistent with that of the corresponding protein. After 100ns, the average values of the four systems were 0.2564, 0.2634, 0.3464 and 0.2502 nm respectively. Except for the system of 600MPa at site 2, the fluctuation of the complex was a little greater than that of the corresponding protein, indicating that the fluctuation of small molecules was greater in the process of binding, which was in agreement with the result reported by Zhan et al (2020) when they study the interaction of β-lactoglobulin with capsaicin. The RMSD of the complex was smaller than that of the corresponding protein, indicating that high pressure treatment could stabilize the binding of small molecules to protein when it bound to protein surface.

**Fig.2.**
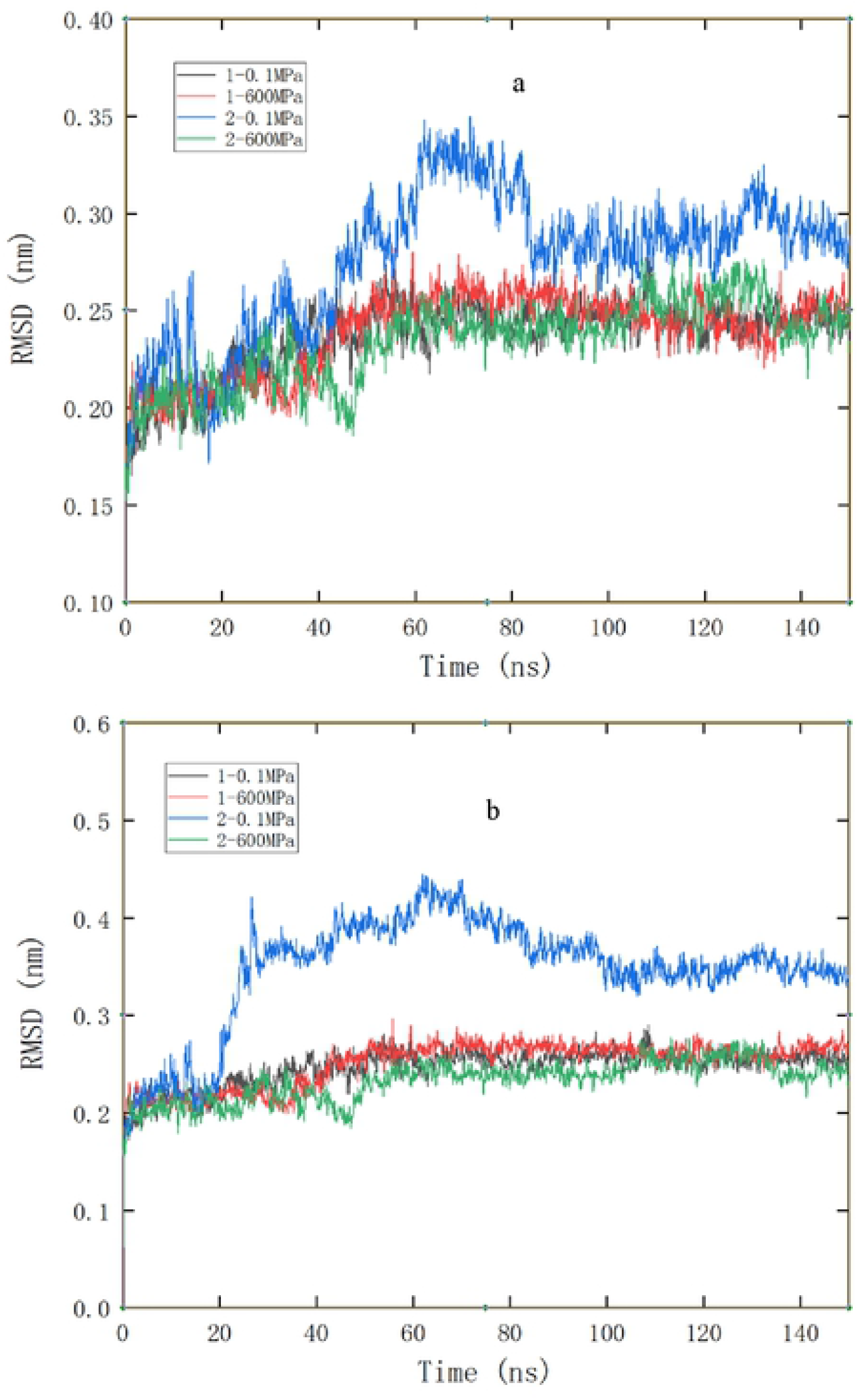
RMSD comparison of protein (a) and complex (b) during different pressure simulation, 1 and 2 represent site 1 and site 2, respectively.

#### 3.2.2 Changes of RMSF in molecular dynamics simulation

RMSF (root mean square fluctuation) reflected the root mean square displacement of each amino acid residue in the protein compared with the average conformation. The greater the fluctuation of the residue, the greater the flexibility of the residue in the pressure treatment. The higher the average flexibility, and the more unstable the protein structure. It can be seen from Fig. 3 that the protein residues fluctuated greatly when the small molecule bound to the protein at site 2 under 0.1 MPa, which was consistent with the RMSD analysis. In terms of the average fluctuation of all residues, the simulation under high pressure was smaller than that under normal pressure, indicating that high pressure could reduce the fluctuation of protein residues. These results were consistent with some other reports, among which Kurpiewska et al (2019) found that the residue displacement of insulin decreased when it was treated at 200 or 500 MPa, and Hata, Nishiyama, and Kitao (2020) also believed that the structural fluctuation of protein decreased under high pressure. In terms of specific amino acid residues, the ones at the entrance of hydrophobic cavity fluctuated greatly (Res110-115, Res85-87), while those near binding site 2 fluctuated slightly in the process of binding.

**Fig.3.**
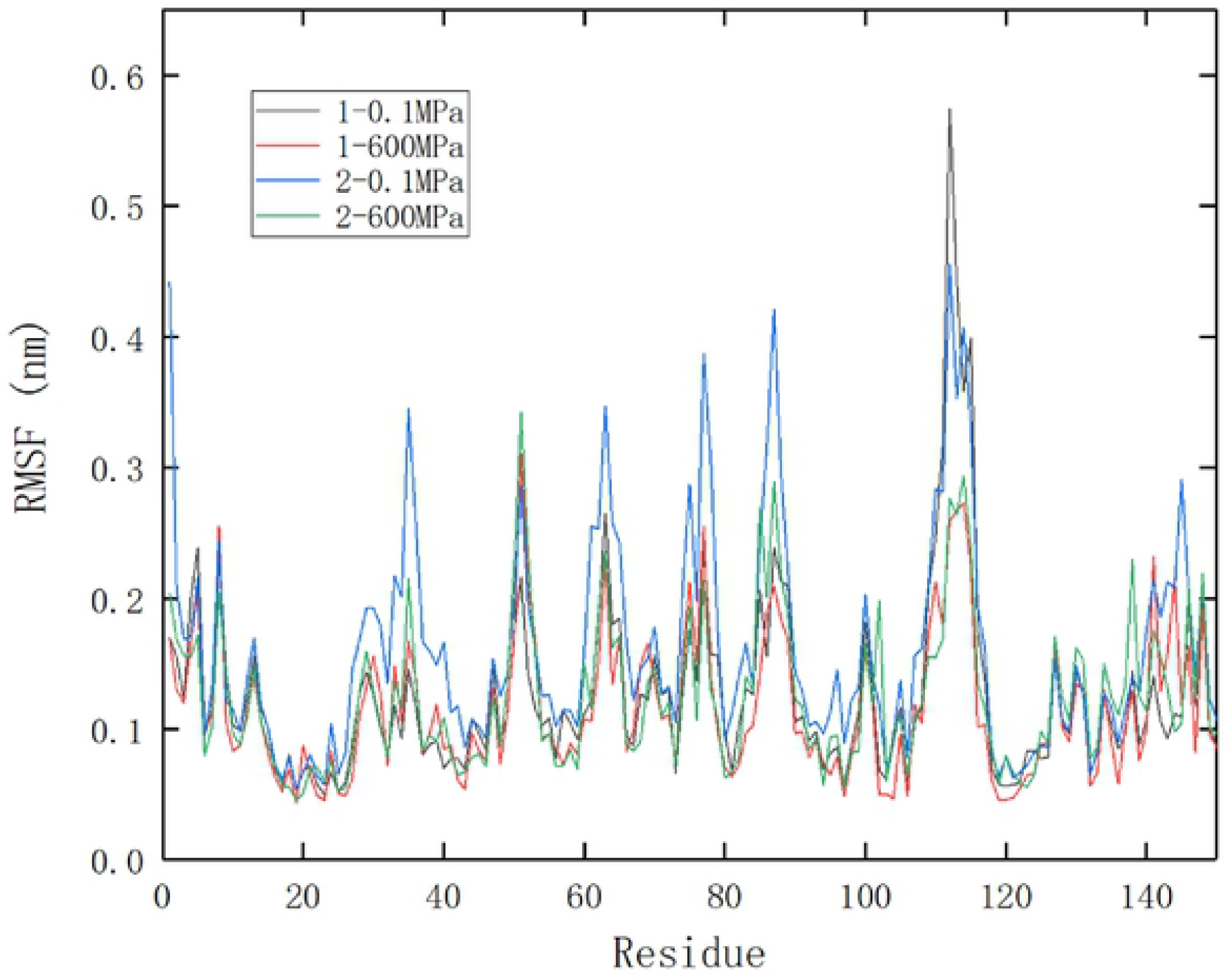
RMSF comparison of protein during different pressure simulation, 1 and 2 represent site 1 and site 2, respectively.

#### 3.2.3 Changes of protein gyration radius in molecular dynamics simulation

Figure 4 showed the change of protein gyration radius (Rg) under different simulation conditions. The smaller the gyration radius was, the more compact the protein structure was. It could be seen that the Rg decreased gradually with the extension of simulation time and remained stable after 70ns. No matter where the small molecule bound, the Rg of protein decreased at 600 MPa, showing that high pressure treatment was conducive to the formation of compact structure of protein, which may be related to the decrease of protein volume under high pressure. According to Le Châtelier principle, the volume of protein decreases under high pressure, which may be due to the deformation of hydrophobic cavity caused by water molecules entering into it under high pressure, resulting in the change of secondary and tertiary structure of protein, and then the volume decreases (Hata et al, 2020).

**Fig.4.**
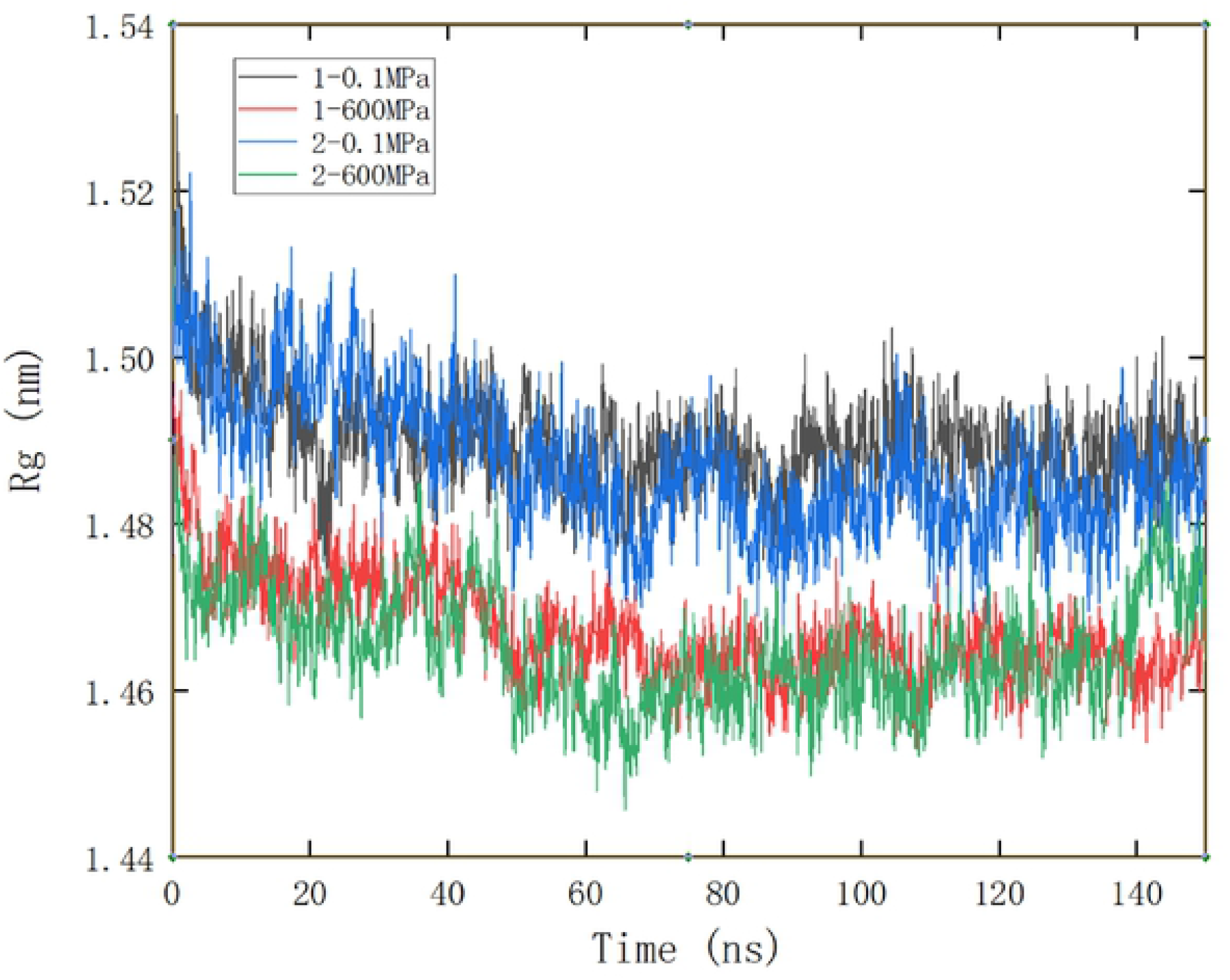
Rg comparison of protein during different pressure simulation, 1 and 2 represent site 1 and site 2, respectively.

#### 3.2.4 Changes of hydrogen bonds between proteins in molecular dynamics simulation

It could be seen from Fig. 5 that after the system was stabilized (after 100ns), the binding of small molecules to protein at site 2 resulted in the decrease of hydrogen bond between proteins, which may be related to the formation of more hydrogen bonds between proteins and small molecules. Hydrogen bond is one of the main non-covalent forces to stabilize protein structure, the drop of hydrogen bond may result in the instability of protein. So the increasing fluctuation of protein RMSD and RMSF when the small molecule bound to protein at site 2 may be related to the decrease of hydrogen bond between protein; however, the fluctuation decreased under 600 MPa, which may be due to the stabilizing effect of pressure. In addition, under high pressure, the hydrogen bonds between protein were stable and the average hydrogen bonds were decreased only by a number of 1.5 compared to 0.1 MPa.

**Fig.5.**
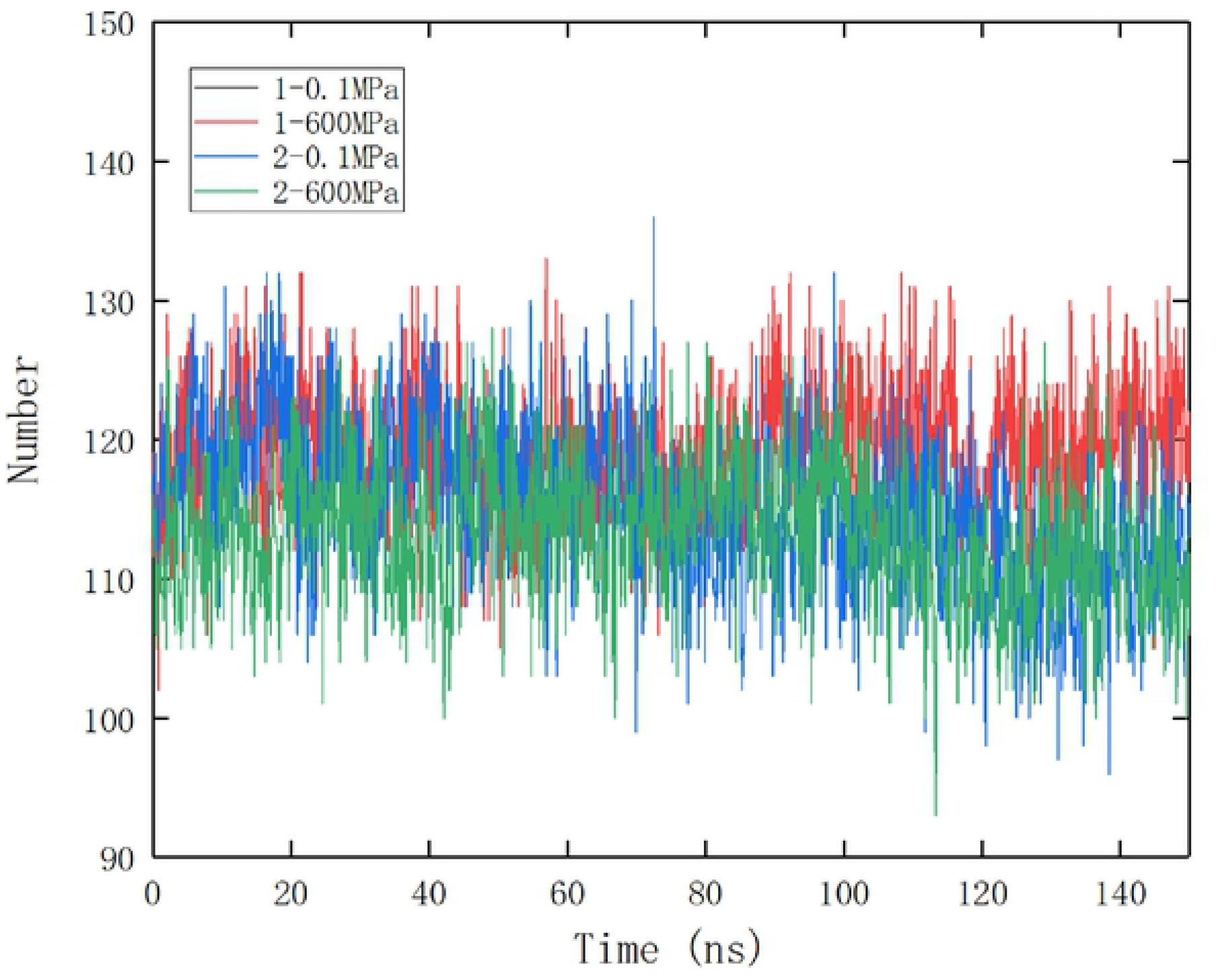
Comparison of hydrogen bond between protein during different pressure simulation, 1 and 2 represent site 1 and site 2, respectively.

#### 3.2.5 Changes of solvent accessible surface area of protein in molecular dynamics simulation

It could be seen from Table 1 that the solvent accessible surface area of protein decreased under 600 MPa, which may be due to the decrease of protein volume. The hydrophilic and hydrophobic areas were reduced simultaneously, but hydrophobic surface area of protein binding EGCG at site 1 decreased more at high pressure. The decrease of hydrophobic surface area may be due to the water molecules infiltrating into the protein molecules under high pressure, so that more hydrophobic areas are exposed to polar aqueous solution (Kolakowski, Dumay, & Cheftel, 2001). This study further showed that when the hydrophobic region bound with small molecules, it was easier to be exposed to polar water solution under high pressure, which may be due to the change of hydrophobic cavity structure caused by the combination of small molecules. Some researchers have also studied the changes of protein hydrophobicity under high pressure by experimental methods, but the results were not consistent, some thought that the hydrophobicity of protein weakened with the increase of pressure (Chen et al, 2019; Hata et al, 2020), while others reported that it increased under high pressure (Qiu, 2014).

#### 3.2.6 Changes of protein secondary structure in molecular dynamics simulation

The secondary structure changes of β-lactoglobulin under different treatment conditions were calculated with DSSP code (Kabsch & Sander, 1983). It could be seen from Table 2 that under 0.1 MPa, the binding of small molecules to protein, no matter at site 1 or site 2, can stabilize the secondary structure of protein with the β-sheet and α-helix increasing while the irregular coil decreasing. Which was in agreement with the result reported by Kanakis et al (2011) when they studied the interaction between milk β-lactoglobulin and tea polyphenols. Under 600 MPa, the secondary structure of the protein was destroyed to some degree, especially the reduction of α–helix. Qiu (2014) also found that the α-helix of myofibrillar protein decreased under high pressure and it was sensitive to pressure. This may be due to the fact that the hydrogen bond of protein was destroyed to a certain extent under high pressure, and the hydrogen bond was crucial to the stability of α-helix. Chen et al (2019) found that the pressure of 400MPa could reduce the α-helix and increase the β-sheet of soybean protein. Chen (2017) reported that the secondary structure of lipase changed little when the pressure was 400MPa, while some α-helix was transformed into irregular coil or β-sheet when the pressure was above 500MPa. These differences may be due to different proteins or different determination methods, most of these reports were measured by experimental methods.

### 3.3 Binding free energy between small molecule and protein using MMPBSA method

It could be seen from Table 3 that the binding between small molecule and protein mainly depended on van der Waals force, followed by electrostatic force and non-polar solvation energy (SASA energy), while the electrostatic interaction was shielded by polar solvation, the values of which were positive and played an unfavorable effect on the binding (Zhan et al, 2020). Under the pressure of 600 MPa, no matter which binding site, the electrostatic energy increased slightly, but the van der Waals and SASA energy decreased compared with the atmospheric pressure simulation system; in addition, the energy consumption of polar solvation increased. Under the combined action of the above forces, the binding free energy decreased significantly at 600 MPa, so the binding of small molecules with protein was not as stable as at normal pressure. From the comparison of two binding sites, the binding energy of small molecules at site 1 was greater than that at site 2 under 0.1 MPa, however, it was just the opposite at 600 MPa, indicating that the protein structure of site 1 was not conducive to the binding of small molecules under high pressure. This was related to a more dramatic decline of van der Waals force at site 1 under 600 MPa; furthermore, a reduction of 4KJ/mol for SASA energy at site 1 were also partially responsible for this change, which closely related to hydrophobic interaction. Chen (2017) discussed the binding energy of lipase with ethyl acetate at 0.1, 200, 400 and 600 MPa and found that the binding energy increased at 200 MPa, but decreased at 400 and 600 MPa, the observed decline of binding energy at 600 MPa was consistent with this study.

The amino acid residues that played a greater contribution (<-0.5KJ/mol) or repulsion (>0.5KJ/mol) to the binding of proteins with small molecules were shown in Fig. 6. It could be seen that at 0.1 MPa, when the small molecule bound at site 1, Leu31, Pro38, Leu39, Val41, Ile71 and Ile84 were the crucial residues contributing to the binding free energy, while residues Lys60, Lys69, Asp85 and Asn90 prevented the protein from binding to small molecule. Under the pressure of 600 MPa, residue Lys60 still prevented the binding of small molecule to protein strongly, and the hindrance of residues Asp85 and Asn90 was greatly reduced, while the hindrance of residue Lys69 was increased. In addition, the role of residues Glu62 and Asn109 changed from contribution to hindrance, and the contribution of residue Leu58 increased significantly under pressure. In Fig. 6 (a), there were 14 hydrophobic amino acids, which contributed −46.93KJ/mol binding energy in total, while in Fig. 6 (b), there were only 8 hydrophobic amino acids with a total energy contribution of −35.61KJ/mol, so it could be judged that the hydrophobic force between small molecule and protein decreased at 600 MPa, which was consistent with the result of SASA energy.

**Fig.6.**
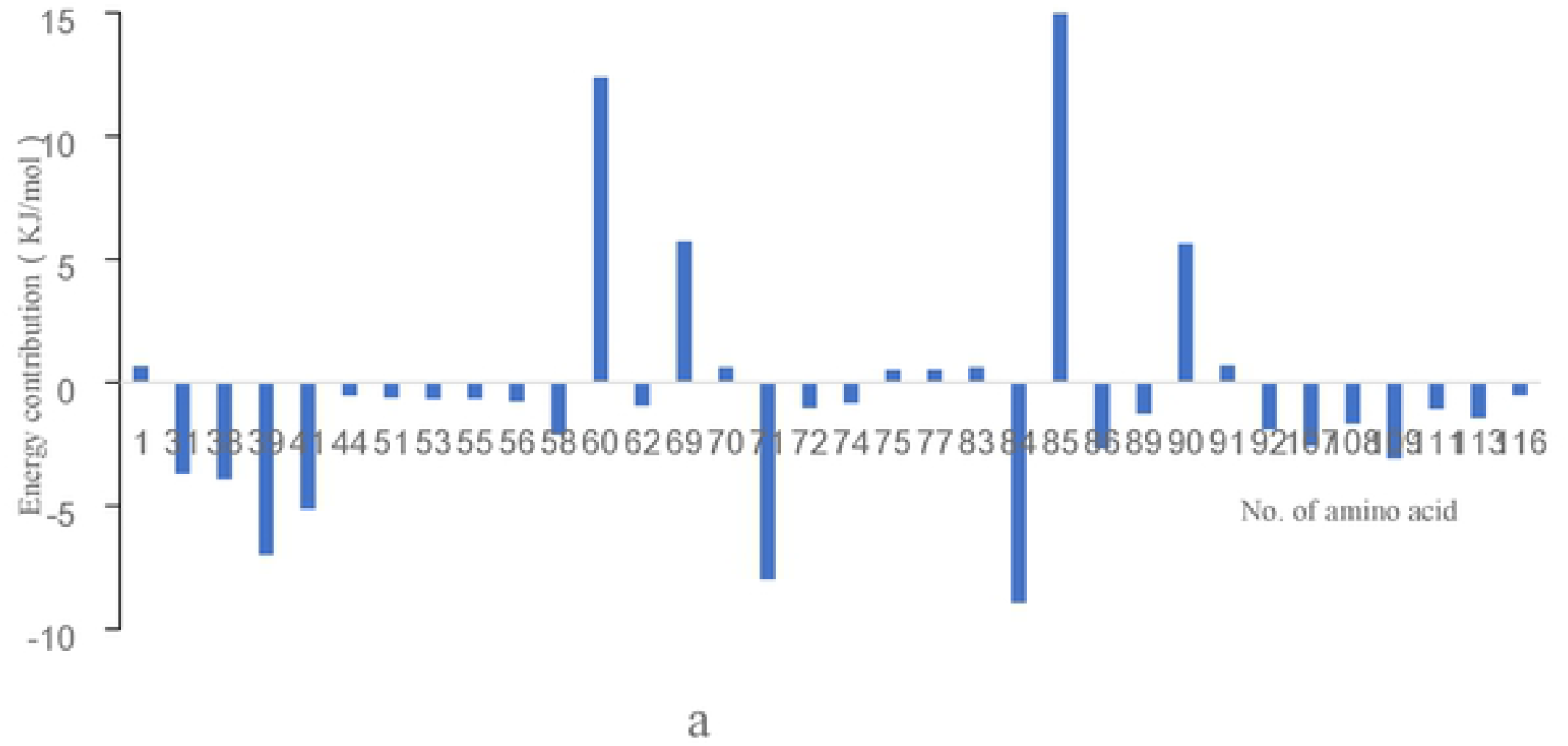
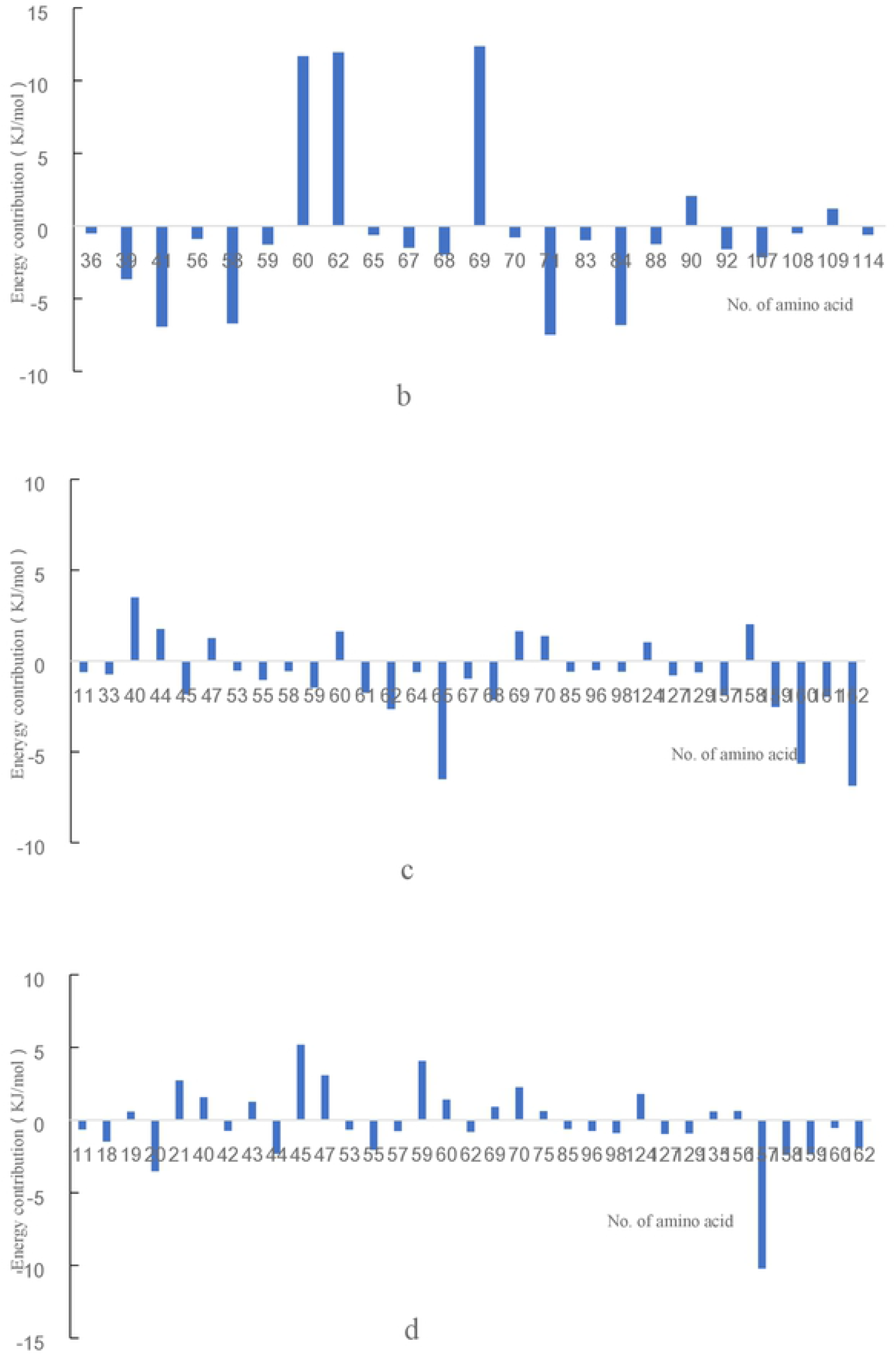
Energy contribution of key amino acid in the combination between protein and EGCG under different treatments (a) site 1-0.l MPa (b)site 1-600MPa (c)site 2-0.lMPa (d)site 2-600MPa.

At atmospheric pressure, when the small molecule bound at site 2, residues Glu65, Cys160 and Ile162 contributed a lot to the binding between EGCG and protein, while residue Glu157 had the greatest role on the interaction under 600 MPa. At 0.1 MPa, there were 8 amino acid residues which had a great hindrance, while the number increased to 14 at 600 MPa, leading to the decrease of total binding free energy. In particular, the contribution of residue Glu65 was significant at atmospheric pressure, but not significant at high pressure. The role of residues Glu45 and Gln59 changed from contribution to hindrance, and the contribution of residues Cys160 and Ile162 also decreased significantly at 600 MPa. In terms of hydrophobic interaction, there were only 4 and 5 hydrophobic amino acids in Fig. 6 (c) and (d), respectively, and their contribution to the total binding free energy was very limited, which may be due to the fact that small molecules bound to the protein surface.

### 3.4 Analysis of possible binding mechanism between protein and small molecule

Figure 7 showed the two-dimensional plot of the interaction between small molecule and protein on site 1 at 0.1MPa (a) and 600MPa (b). It could be seen that at 0.1 MPa, the small molecule formed four hydrogen bonds with protein, two with Asp85, and one with Asn109 and Glu112 respectively. At 600 MPa, five hydrogen bonds were observed, including two with Glu62, and one with Leu58, Ala67 and Pro38 respectively. It could be seen that the hydrogen bonds formed under normal and high pressure were completely different, indicating that the binding of small molecules with protein changed greatly under high pressure. Under both pressures, the small molecule interacted hydrophobically with the 12 amino acids of the protein, but the hydrophobic environment changed a lot, at normal pressure, 10 of 12 amino acids belonged to hydrophobic amino acids, while at high pressure, only 6 hydrophobic amino acids. These results were also consistent with the conclusion in 2.3 that the hydrophobic interaction was weakened under high pressure when small molecule bound at position 1 of protein. So, the binding law and environment of small molecule and protein changed greatly at 600 MPa, and thus resulting in the change of binding energy.

**Fig.7.**
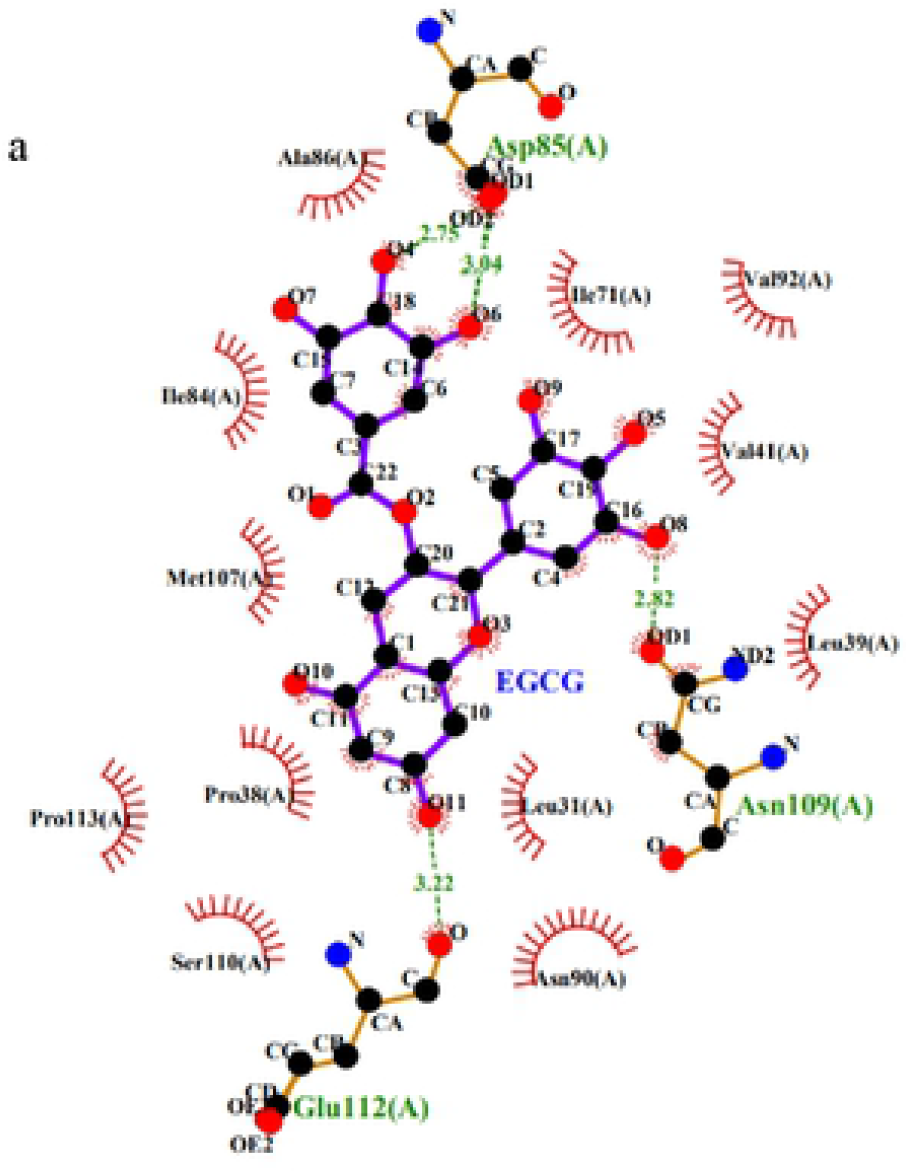
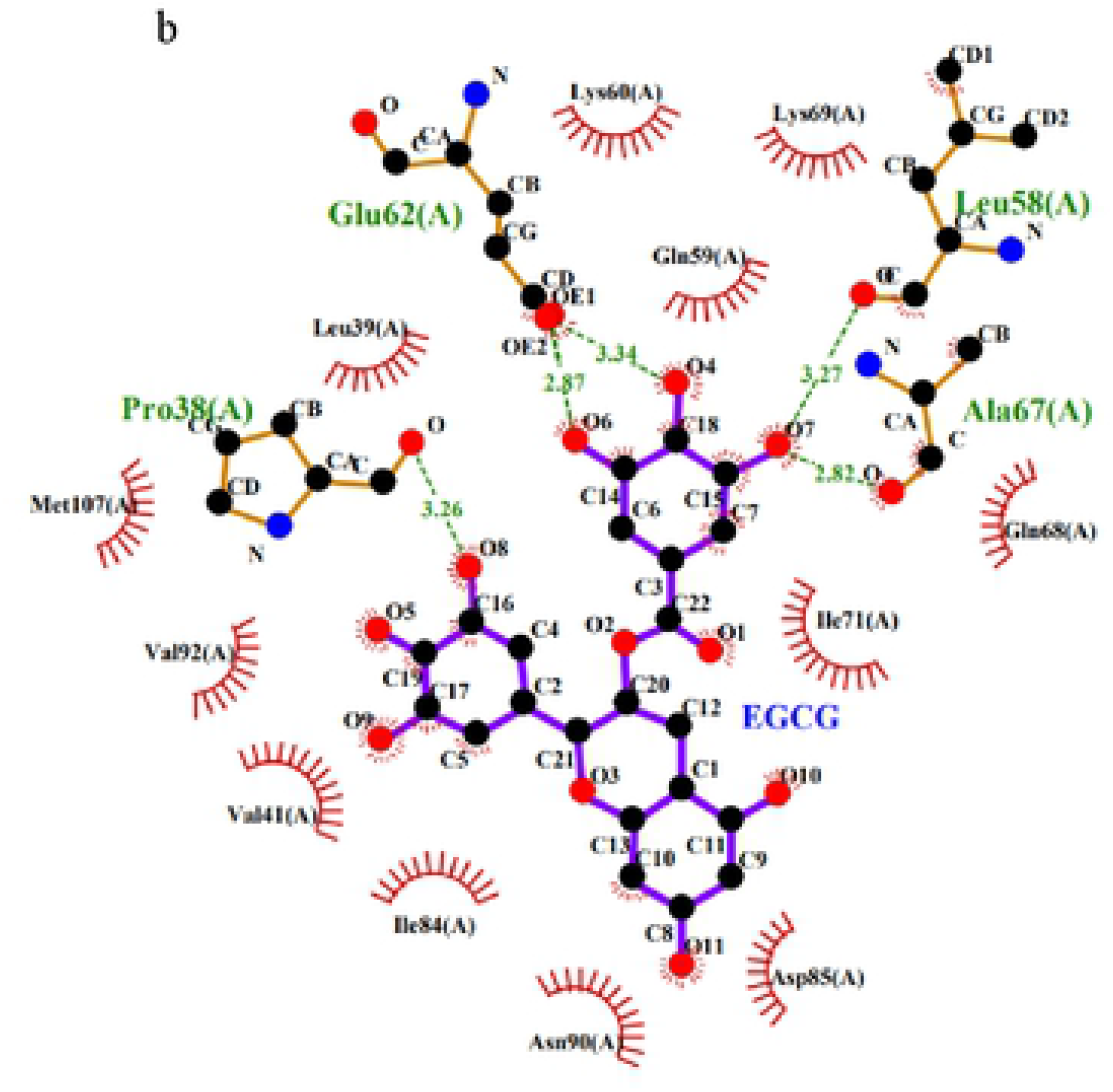
The 2-D plot for interaction between EGCG and β-lactoglobulin at site 1 under 0.1 MPa (a) and 600 MPa (b)

Figure 8 showed the two-dimensional plot of the interaction between small molecule and protein on site 2 at 0.1MPa (a) and 600MPa (b). It could be seen that at 0.1 MPa, the EGCG formed six hydrogen bonds with protein, two with Glu44, and one with Gln59, Glu68, Glu158, and Cys160 respectively. At 600 MPa, seven hydrogen bonds were found, including two with Glu158, and one with Trp19, Glu44, Glu45, Gln159, and Gln159 respectively. There were three same hydrogen bonds at different pressure. More hydrogen bonds were observed between small molecule and protein at position 2 than at position 1, which may be related to the decrease of hydrogen bonds between proteins when EGCG binding to the surface of protein. Site 2 was located on the surface of protein, so the hydrophobic interaction between small molecule and amino acid residues was weak, and there were only three and four residues interacted hydrophobically with EGCG at 0.1 and 600 MPa respectively.

**Fig.8.**
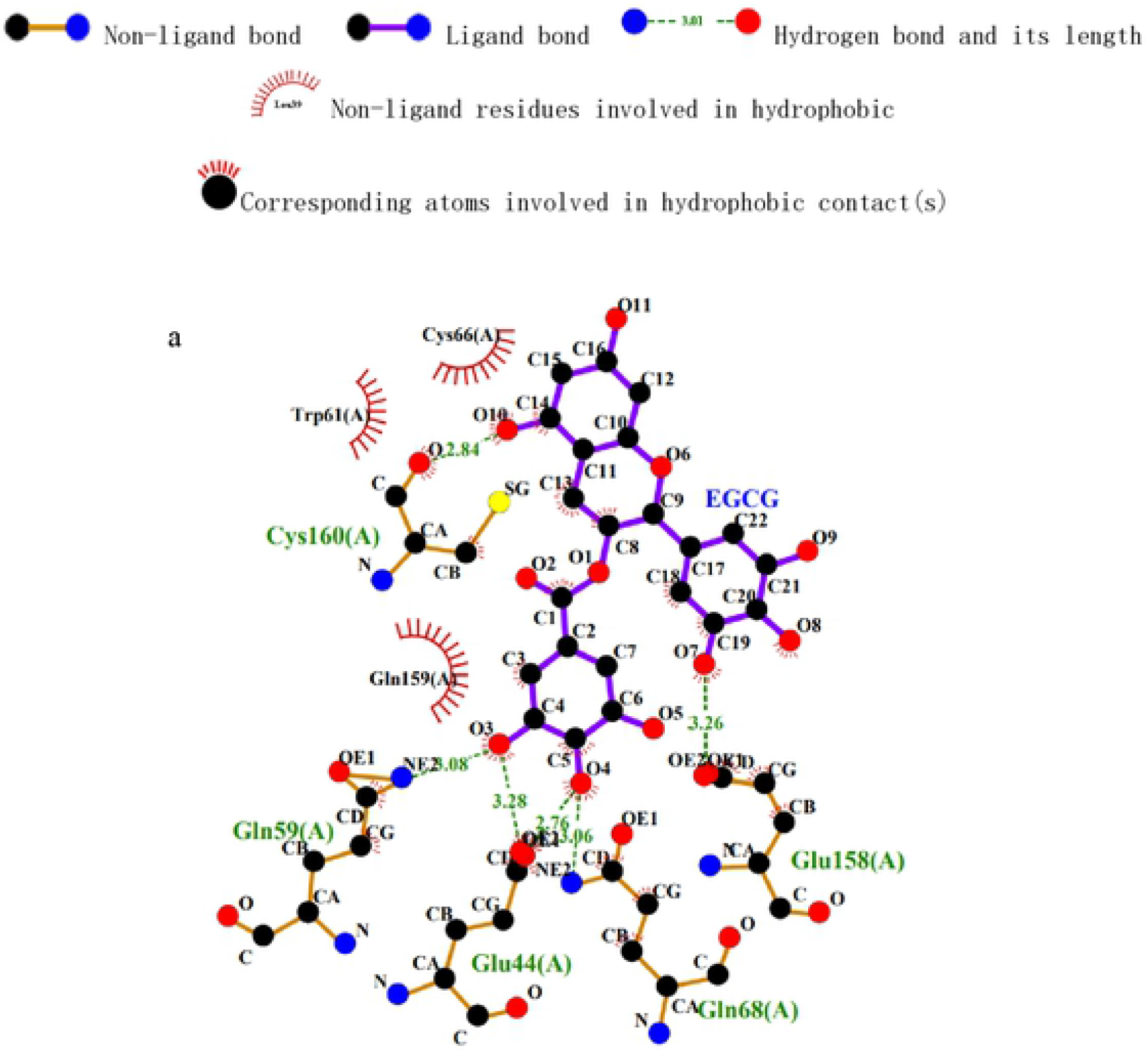
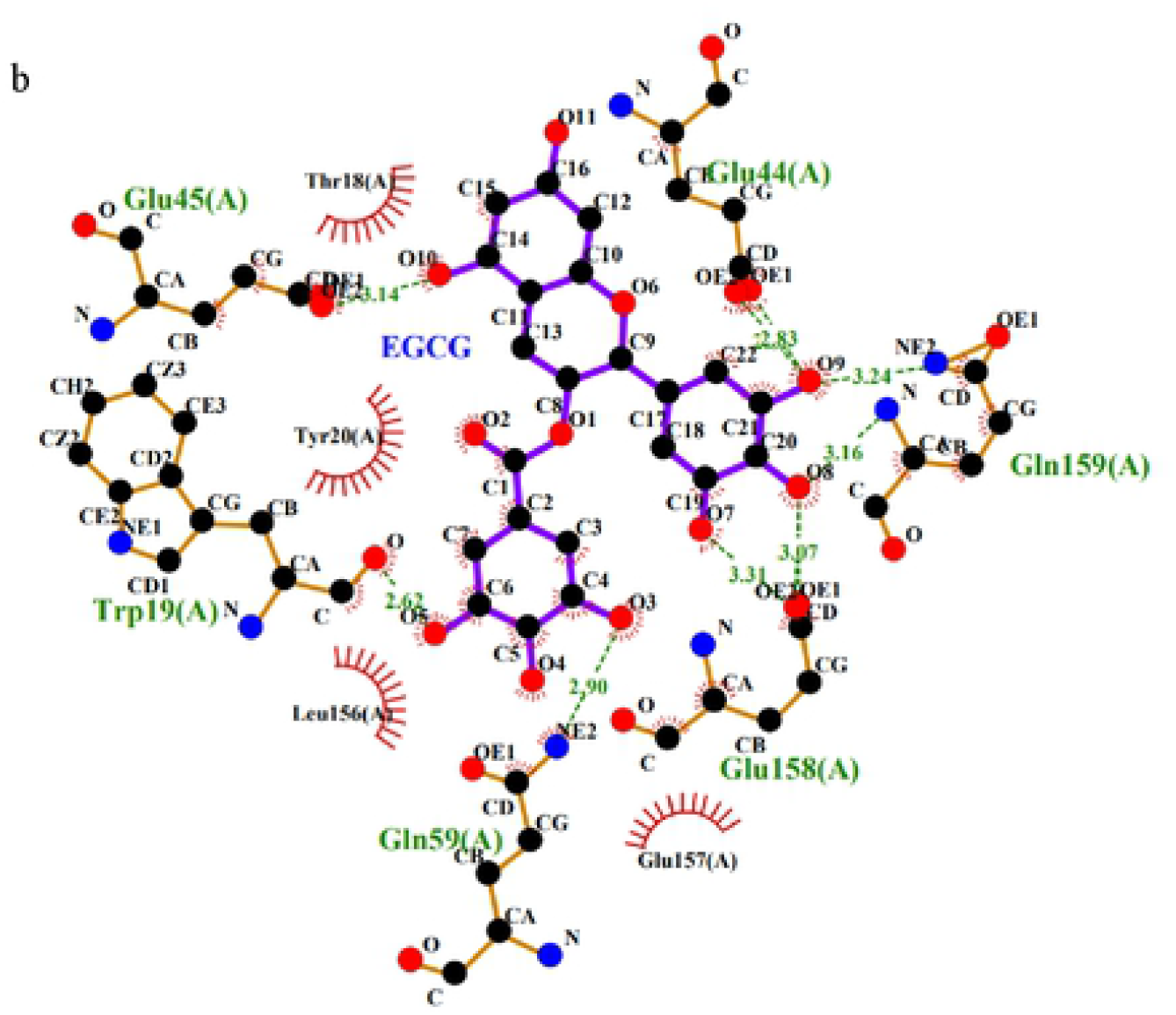
The 2-D plot for interaction between EGCG and β-lactoglobulin at site 2 under 0.1 MPa (a) and 600 MPa (b)

So, there existed van der Waals force, hydrophobic force, and hydrogen bond simultaneously when small molecule bound in hydrophobic cavity, while hydrophobic force was very small when EGCG bound on the surface of protein, mainly van der Waals force and hydrogen bond. At 600 MPa, the reduction of binding free energy at site 1 was due to the decrease of hydrophobicity and van der Waals force, while the reduction of van der Waals force was the main reason for site 2.

In order to further understand the effect of high pressure on the surface structure of protein and its binding to small molecules, PyMOL software was used to draw the surface structure diagram of protein molecule. Fig. 9 (a) and (b) showed the average surface structure of protein and the binding of polyphenols to protein at site 1 during 100-150 ns of simulation at 0.1 and 600 MPa, respectively. It could be seen that the binding site of the small molecule remained at the top of the hydrophobic cavity under 600 MPa, however, the binding posture of small molecules changed greatly, which was consistent with the results of the previous analysis on hydrogen bond and hydrophobic environment. In addition, the surface structure of hydrophobic cavity changed obviously under high pressure, which may also affect the binding of small molecules to protein. The average surface structure of protein and the binding of polyphenols to protein at site 2 during 100-150 ns of simulation at 0.1 and 600 MPa were shown in Fig. 9 (c) and (d), respectively. It could be seen that under high-pressure, the small molecule had a large displacement on the protein surface, which may be one of the main reasons for the great changes in binding energy at 600 MPa.

**Fig.9.**
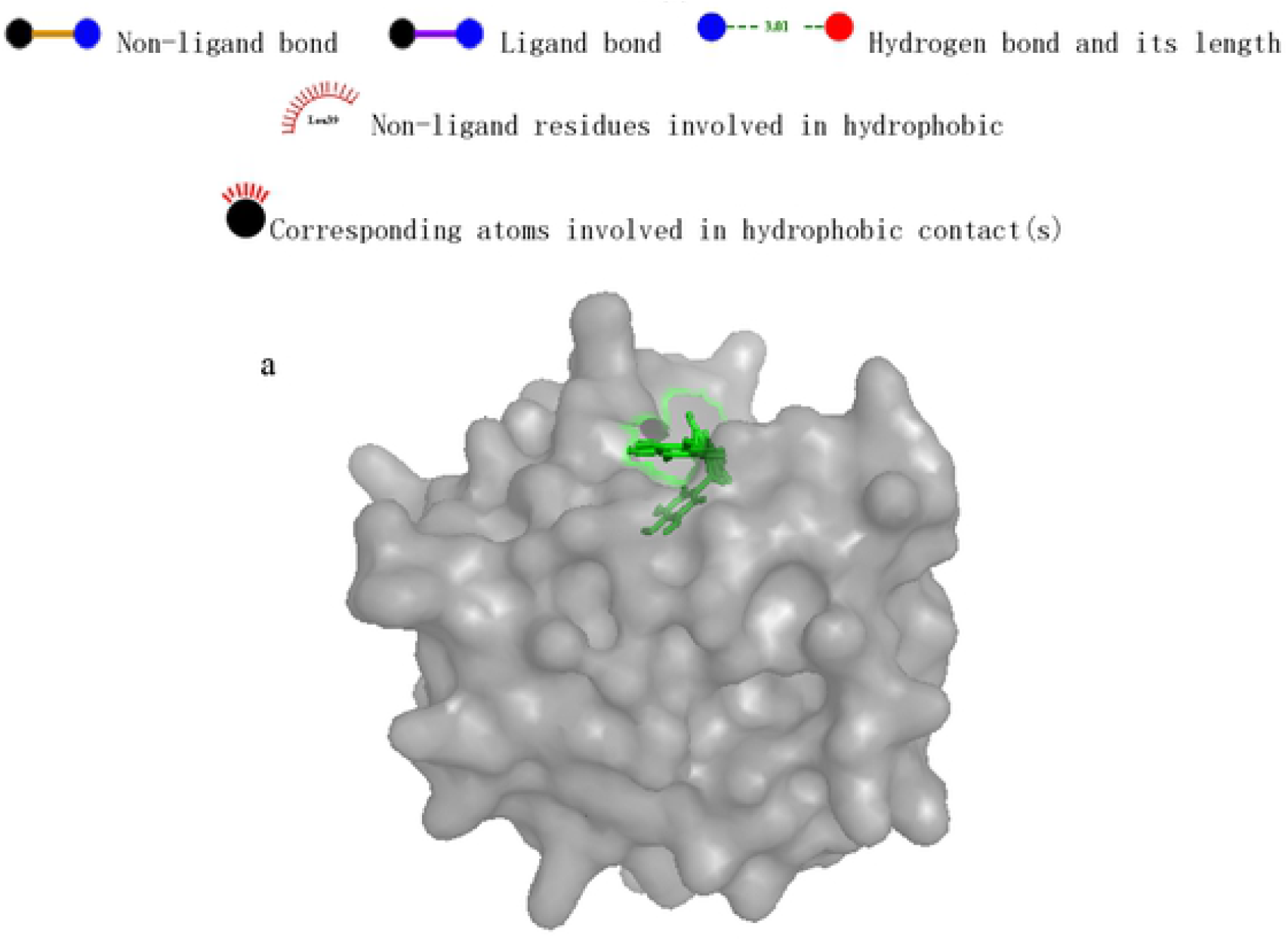
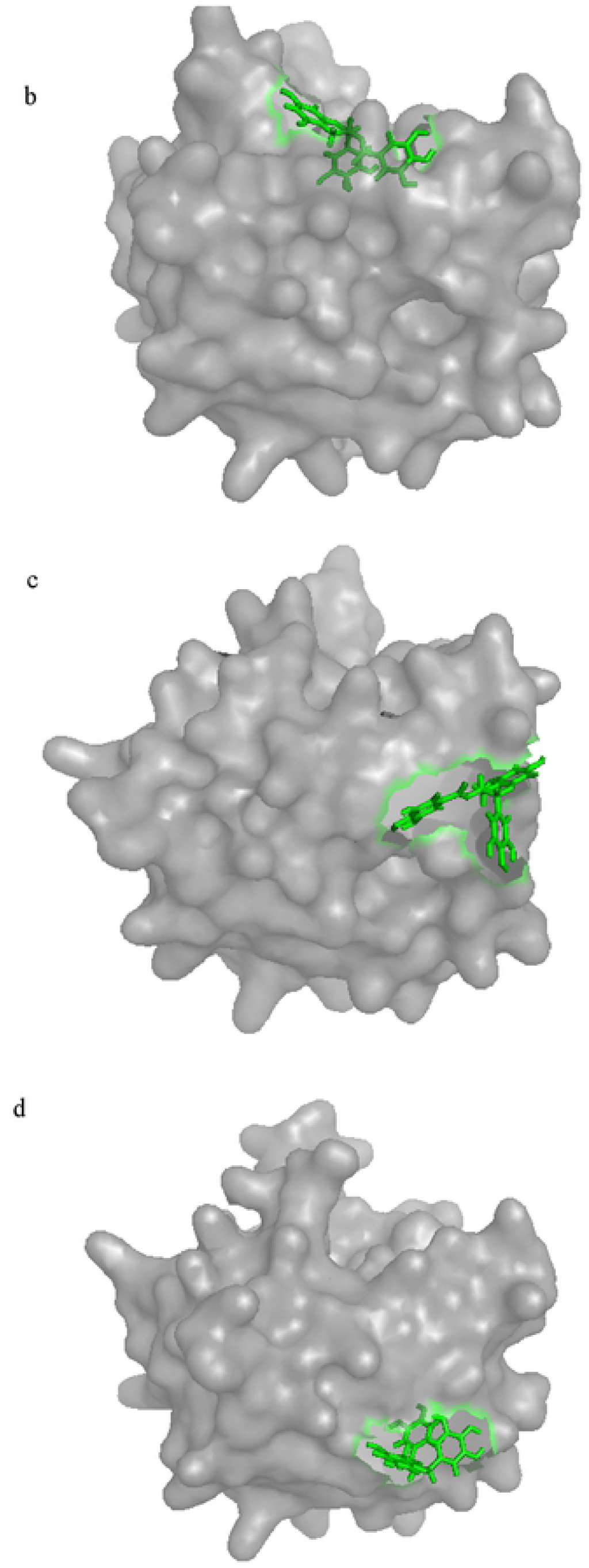
Surface structure and polyphenol binding in protein under 0.1 (a) and 600 (b) MPa at site 1, and under 0.1 (c) and 600 (d) MPa at site 2, respectively.

## Conclusion

EGCG interacted with β-lactoglobulin mainly in two sites. The binding at hydrophobic cavity (site 1) mainly depended on van der Waals force, hydrophobic interaction and hydrogen bond. Under pressure of 600 MPa, the binding free energy significantly reduced mainly due to the van der Waals force and hydrophobic interaction. The interaction at protein surface (site 2) mainly depended on van der Waals force and hydrogen bond. Van der Waals force was far less than while hydrogen bond interaction was stronger than those at site I, in conjunction with the fact that weaker hydrophobic interaction in site 2, the total binding free energy of small molecules at this site was lower than that in hydrophobic cavity at 0.1 MPa. Under the pressure of 600 MPa, the binding site of small molecules on the protein surface shifted significantly, and the van der Waals force decreased significantly, resulting in the binding free energy also decreased significantly, but higher than that of site 1 under 600 MPa. Therefore, the best binding site of EGCG was in the hydrophobic cavity at atmospheric pressure, while the binding free energy in protein surface was higher at 600 MPa. The findings in this study laid a good theoretical foundation for further improving the quality of tea milk beverage and the application of high-pressure technology in milk beverage.

## Acknowledgments

This work was supported by National Key Research and Development Plan (2018YFD0502404).

